# Separate Gut Plasma Cell Populations Produce Autoantibodies Against Transglutaminase 2 and Transglutaminase 3 in Dermatitis Herpetiformis

**DOI:** 10.1101/2023.05.31.542741

**Authors:** Saykat Das, Jorunn Stamnaes, Esko Kemppainen, Kaisa Hervonen, Knut E.A. Lundin, Naveen Parmar, Frode L. Jahnsen, Jørgen Jahnsen, Katri Lindfors, Teea Salmi, Rasmus Iversen, Ludvig M. Sollid

## Abstract

Dermatitis herpetiformis (DH) is an inflammatory skin disorder often considered as an extraintestinal manifestation of celiac disease (CeD). Hallmarks of CeD and DH are autoantibodies to transglutaminase 2 (TG2) and transglutaminase 3 (TG3), respectively. DH patients have autoantibodies reactive with both transglutaminase enzymes. We here report that in DH both gut plasma cells and serum autoantibodies are specific for either TG2 or TG3 with no TG2-TG3 cross-reactivity. By generating monoclonal antibodies from TG3-specific duodenal plasma cells of DH patients, we define three conformational epitope groups. Both TG2-specific and TG3-specific gut plasma cells have few immunoglobulin mutations, and the two transglutaminase-reactive populations show distinct selection of certain heavy and light chain V-genes. Mass spectrometry analysis of TG3-specific serum IgA corroborates preferential usage of *IGHV2-5* in combination with *IGKV4-1*. Collectively, our results demonstrate parallel induction of anti-TG2 and anti-TG3 autoantibody responses involving separate B-cell populations in DH patients.

## Introduction

Dermatitis herpetiformis (DH) typically manifests by an itchy, blistering rash on the extensor surfaces of knees, elbows, and buttocks.^[1]^ The disease diagnosis is made based on detection of IgA deposits in the papillary dermis of unaffected skin.^[2]^ DH is related to celiac disease (CeD), as both conditions depend on dietary exposure to cereal gluten proteins.^[3]^ Both diseases demonstrate the same strong association to the HLA-DQ allotypes DQ2.5, DQ2.2 and DQ8, and both types of patients have CD4+ T cells specific for deamidated gluten peptides.^[4]^ In both diseases, the immune response to gluten drives the generation of autoantibodies to transglutaminase enzymes; in CeD to transglutaminase 2 (TG2) and in DH to transglutaminase 3 (TG3) as well as to TG2.^[5]^ Transglutaminases are a family of enzymes characterized by Ca2+ dependent and sequence-specific targeting of polypeptide glutamine residues, leading either to cross-linking via attachment of the side chain to a primary amine (such as a lysine residue; transamidation) or conversion into glutamic acid through hydrolysis (deamidation).^[6]^ Gluten peptides that have undergone transglutaminase-mediated deamidation bind better to the HLA-DQ2.5, HLA-DQ2.2 and HLA-DQ8 molecules, thus explaining why CD4+ T cells of CeD and DH patients recognize such post-translationally modified peptides.^[7]^ While both TG2 and TG3 are able to generate the immunodominant gluten T-cell epitopes, their fine specificities are slightly different.^[8]^

Serum anti-TG2-IgA antibody tests are widely employed for diagnosis of CeD due to their excellent sensitivity and specificity.^[9]^ Moreover, IgA and IgM plasma cells specific for TG2 are abundant as effector B cells in the small intestine lamina propria of CeD patients.^[10]^ To explain the connection between gluten exposure and the formation of TG2-specific autoantibodies in CeD, it has been proposed that TG2-specific B cells take up TG2-gluten enzyme-substrate complexes via their B-cell receptor (BCR), thereby allowing presentation of deamidated peptides to gluten-specific CD4+ T cells. The T cells then provide activation signals to the B cells, leading to production of TG2-specific autoantibodies.^[11]^ TG2-specific gut plasma cells exhibit preferential usage of certain immunoglobulin (Ig) heavy and light chain variable region gene segments,^[10]^ and this selection was shown to reflect targeting of particular N-terminal TG2 epitopes.^[12]^ Among TG2-specific antibodies, the combination of an *IGHV5-51* heavy chain and an *IGKV1-5* light chain is particularly prominent and clonally expanded.^[13]^ B cells expressing such BCRs were able to efficiently present deamidated gluten peptides to CD4+ T cells through favorable binding of TG2-gluten enzyme-substrate complexes, potentially explaining the V-gene bias.^[14]^ Whether a similar hapten-carrier-like mechanism is involved in gluten-dependent production of TG3-specific autoantibodies in DH is not known, but recently it was reported that DH patients have TG3-specific plasma cells in the small intestine lamina propria.^[15]^ A previous study on TG2- and TG3-binding serum antibodies indicated that there might exist cross-reactive antibodies capable of binding both enzymes.^[5]^ Due to their structural similarity and because DH usually develops as a late complication of CeD, it has been postulated that the response against TG3 could arise from an initial TG2-focused response.^[16]^

To gain insight into the origin of TG3-specific autoantibodies and understand their connection to the anti-TG2 response, we here characterize TG3-binding autoantibodies in DH patients by assessing serum antibody reactivity and by generation of monoclonal antibodies (mAbs) from TG3-binding duodenal plasma cells. We demonstrate that TG3-binding antibodies are highly specific and do not show cross-reactivity to TG2. Both TG2- and TG3-binding antibodies are produced in DH with an apparent skewing toward production of anti-TG2 IgA in most patients. The two transglutaminase-specific responses show similar features, including few Ig mutations and preferential usage of particular V-gene segments. Based on our results, we propose that anti-TG2 and anti-TG3 autoantibodies in DH are generated through parallel mechanisms, but with involvement of separate antigen-specific populations of B cells.

## Results

### DH patients have specific serum antibodies to TG2 and TG3

To compare the anti-transglutaminase responses in patients with untreated CeD and DH, we assessed binding of serum IgA and IgG to TG2 or TG3 by ELISA (Figure 1A-B). Whereas CeD and DH patients both showed binding to TG2, antibody binding to TG3 was specific to DH. For both TG2 and TG3, the IgA antibodies were more accurate disease markers than IgG, as not all patients showed increased levels of TG2- and TG3-specific IgG compared to healthy donors. Hence, both TG2- and TG3-specific serum antibodies likely derive from mucosal immune reactions. There was poor correlation between the levels of anti-TG2 and anti-TG3 serum IgA in DH patients, suggesting that the two types of autoantibodies involve separate populations of plasma cells (Figure 1C).

**Figure 1.**
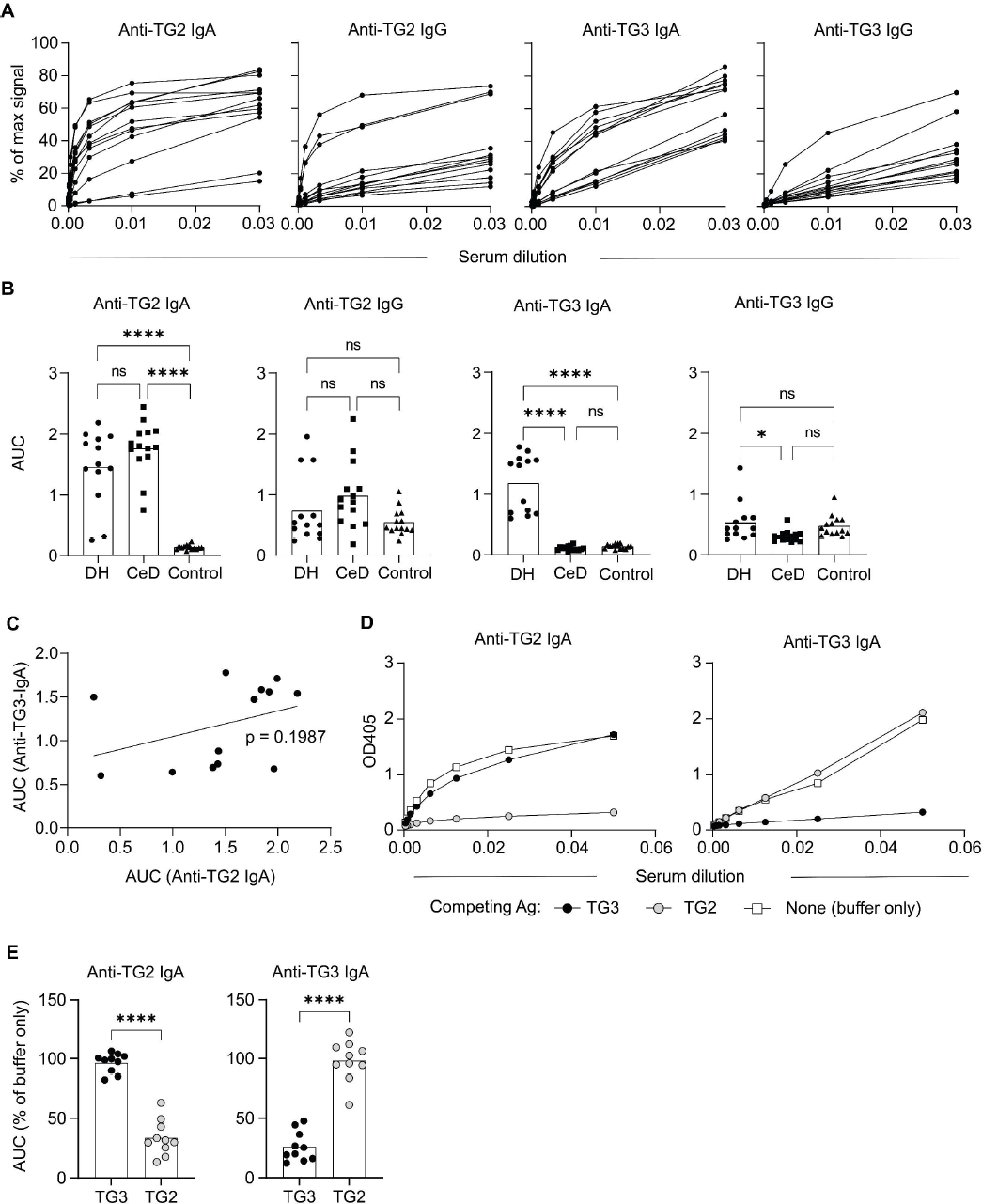
Serum IgA and IgG antibodies reactive to TG3 and TG2. (A) Titration binding curves showing TG2 and TG3 reactivity of serum IgA and IgG in untreated DH patients as assessed by ELISA. Signals are given relative to the maximum signals obtained with reference IgA and IgG mAbs. (B) Comparison of antibody levels in DH patients (*n*=13), CeD patients (*n*=14) and control donors (*n*=14). Signals are given as area-under-curve (AUC) values obtained from ELISA binding curves as the ones shown in (A). Difference between groups was evaluated by one-way ANOVA with Holm-Sidak multiple comparisons correction. (C) Correlation between the levels of anti-TG2 IgA and anti-TG3 IgA in DH patients. The p-value was calculated by Pearson correlation analysis. (D-E) Assessment of serum IgA cross-reactivity between TG2 and TG3. (D) Representative ELISA binding curves showing reactivity of serum IgA from a single DH patient to TG2 or TG3 with or without pre-incubation with the indicated immobilized antigens (Ag). (E) AUC values obtained from binding curves as the ones shown in (D) summarizing results from 10 DH patients. Statistical difference between TG3 and TG2 preincubation was analyzed by a paired t-test. ****p<0.0001, **p<0.01, *p<0.05

Previous studies reported that DH patients have TG3-reactive autoantibodies that also bind to TG2.^[5, 16]^ To test for potential cross-reactivity, we incubated serum samples of DH patients with immobilized TG2 or TG3 and assessed remaining IgA reactivity in the supernatant (Figure 1D-E). Preincubation with either TG2 or TG3 abolished binding to the same transglutaminase antigen, but did not affect binding to the other one. Only in one out of ten patients did we observe a substantial (39%) drop in TG3 reactivity upon incubation with TG2. Thus, our results indicate that DH patients have separate serum antibody populations targeting TG2 and TG3 with no or very limited cross-reactivity.

### TG3-specific plasma cells are present in duodenal biopsies of DH patients

To investigate whether we could identify plasma cell populations mirroring the TG2- and TG3-specific serum antibodies, we co-stained duodenal biopsy single-cell suspensions of CeD or DH patients with TG2 and TG3 and detected reactive cells by flow cytometry (Figure 2A-B and Figure S1). In agreement with previous findings,^[13, 17]^ a population of TG2-binding plasma cells comprising, on average, 14% of all IgA plasma cells could be detected in untreated CeD patients (range, 3.8-24.3%). A similar population of cells was present in DH patients (range, 0.8-25.5%). In addition, DH patients had a population of TG3-binding plasma cells that was not present in CeD patients. Notably, we did not detect any double-positive cells, indicating that gut plasma cells show strict specificity to either TG2 or TG3. Although there was substantial variation, the frequency of TG3-specific plasma cells in DH patients was, on average, ten times lower than that of TG2-specific plasma cells (mean: 1.4% vs 13.9%). One patient had few TG2-specific plasma cells (0.8%) and also low serum anti-TG2 IgA (EC50 = 1:7), but a high frequency of TG3-specific gut plasma cells (2.6%). Further, the ratio between TG2- and TG3-specific gut plasma cells reflected the relative levels of serum IgA against the two enzymes (Figure 2C). Thus, the plasma cell populations we detected in duodenal biopsies appear to be directly related to the serum IgA responses. Similar to what has previously been reported for TG2-specific plasma cells in CeD, the majority of the TG3-specific cells in DH were CD19+, indicating that they are newly generated and result from ongoing immune reactions (Figure 2D).^[17-18]^

**Figure 2.**
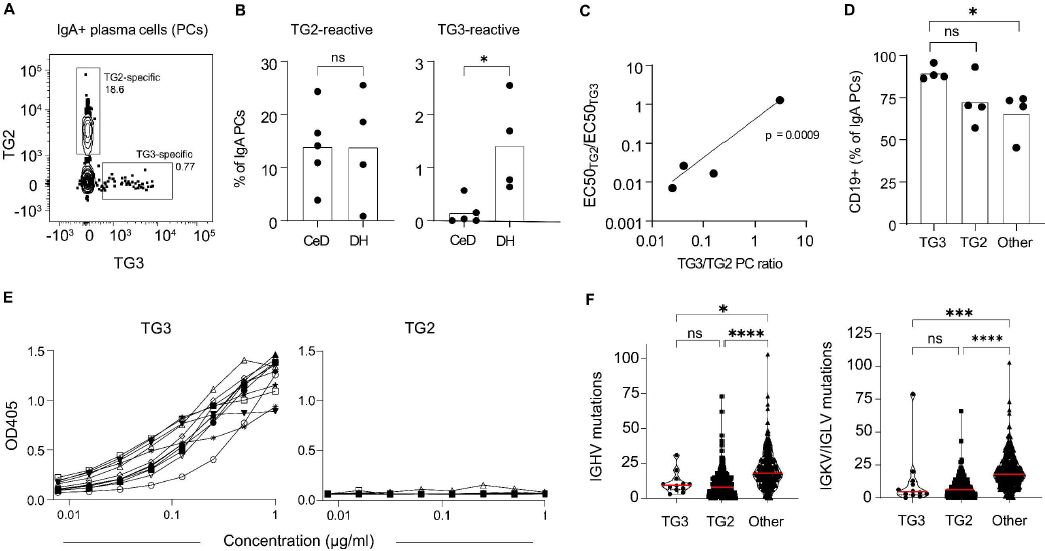
TG2- and TG3-binding plasma cells in duodenal biopsies of DH and CeD patients. (A) Representative flow cytometry plot showing staining of IgA gut plasma cells of a DH patient with TG2 and TG3. (B) Percentage of TG2- and TG3-binding IgA plasma cells in gut biopsies of CeD (*n*=5) and DH (*n*=4) patients. Statistical difference was evaluated by an unpaired t-test. (C) Correlation between the ratio of TG2- and TG3-specific gut plasma cells (PCs) and the relative levels of TG2- and TG3-specific serum IgA given as EC50 values determined by ELISA in four DH patients. The p-value was calculated by Pearson correlation analysis. (D) Expression of CD19 among TG3-specific, TG2-specific and non-TG3-non-TG2-specific (other) IgA plasma cells in duodenal biopsies of DH patients. Statistical difference was evaluated by paired a t-test. (E) ELISA binding curves showing TG3 and TG2 reactivity of recombinant mAbs (*n*=12) generated from TG3-binding gut plasma cells of DH patients. (F) Number of *IGHV* and *IGKV/IGLV* mutations among the TG3-specific mAbs. The numbers were compared to previously reported sequences obtained from TG2-specific (*n*=286) and non-TG2-specific (*n*=338) gut plasma cells isolated from CeD patients.^[13-14]^ Red lines indicate medians, and difference between groups was evaluated by a Kruskal-Wallis H test with Dunn’s multiple comparisons correction. ****p<0.0001, ***p<0.001, *p<0.05

Since our results indicate that the IgA responses against TG2 and TG3 are induced in the gut, we assessed expression of both antigens by immunofluorescence staining of frozen intestinal tissue sections. Whereas TG3 is only expressed in the crypt compartment in the small intestine with little or no expression in the colon, TG2 is abundantly expressed both in the small intestine and the colon (Figure S2A). These results were corroborated by analysis of a published proteome data set of CeD small intestine,^[19]^ which revealed higher expression of TG2 than TG3 (Figure S2B). As expected, TG3 expression could also be detected in the epidermis of the skin using the same immunofluorescence staining procedure (Figure S2C). Our results confirm that both TG2 and TG3 are present in the small intestine and presumably capable of driving autoantibody production.

To further characterize the anti-TG3 response in DH, we generated 12 recombinant mAbs from single TG3-binding IgA gut plasma cells of three individual DH patients (Table S1). In agreement with the gut plasma cell staining pattern and serum antibody reactivity, the mAbs showed specific binding to TG3 without any detectable cross-reactivity to TG2 (Figure 2E). The mAbs displayed similar levels of Ig mutations as what has previously been reported for TG2-specific plasma cells in CeD (Figure 2F).^[10, 13]^ Thus, both TG2-specific and TG3-specific plasma cells accumulate fewer mutations than other IgA gut plasma cells, indicating that they are generated through equivalent mechanisms, possibly outside of germinal centers.^[7]^ Among the 12 TG3-specific mAbs, four used *IGHV2-5* in combination with *IGKV4-1* (Table S1). Three of these were clonally related cells, while the last one was isolated from a different patient. Although our data include few sequences, this observation could indicate that the anti-TG3 response in DH selects for specific combinations of heavy and light chain V-gene segments, similar to what has been described for TG2-specific antibodies in CeD.^[13]^

### TG3-specific mAbs target distinct conformational epitope groups

TG2-specific antibodies in CeD show preferential targeting of particular epitopes.^[12]^ To address if TG3-specific antibodies in DH are also directed toward common epitopes, we assessed the degree of competition between individual TG3-specific mAbs as assessed by competition ELISA (Figure 3A). Based on the ability of the mAbs to compete with each other for binding, we define three epitope groups, where each group reflects a discrete region of the enzyme targeted by a subset of mAbs. The epitopes recognized by the mAbs are strictly conformational, as all antibody reactivity was lost upon denaturation of TG3 (Figure 3B).

**Figure 3.**
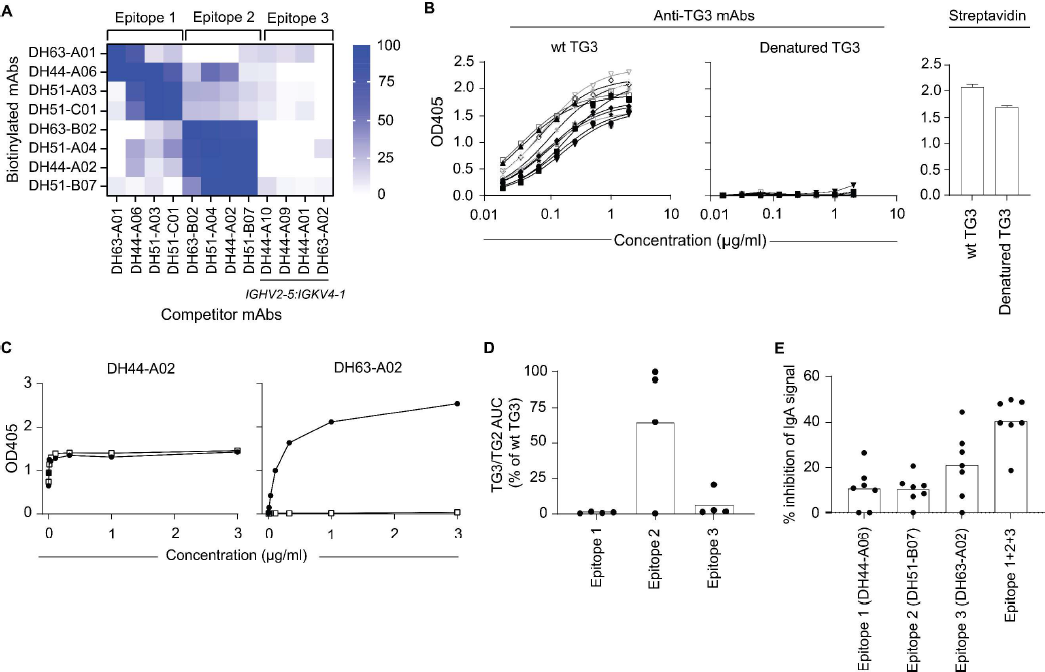
TG3-specific mAbs target three distinct conformational epitope groups. (A) Heatmap showing the degree of competition between individual mAbs for binding to TG3 in ELISA. In each case, the ability of an unlabeled competitor to inhibit binding of a biotinylated mAb was assessed. Based on the reactivity pattern, mAbs were divided into epitope groups 1-3. The four mAbs using *IGHV2-5:IGKV4-1* (epitope group 3) lost reactivity upon biotinylation and were therefore only used as competitors in the assay. Although not directly assessed, these mAbs likely bind to the same epitope. (B) Titration binding curves showing reactivity of the mAbs to wild-type (wt) and denatured TG3 in ELISA. Detection of the biotinylated antigens with streptavidin showed approximately equal levels of coating for wt and denatured TG3. Error bars indicate SD based on eight individual measurements. (C) Examples of ELISA binding curves for mAbs that either bind (DH44-A02) or do not bind (DH63-A02) to a TG3/TG2 chimeric protein. (D) Summary data for all 12 anti-TG3 mAbs showing binding to the TG3/TG2 chimera. Levels are given as area-under-curve (AUC) values obtained from binding curves as the ones shown in (C). (E) Binding of serum IgA of DH patients (*n*=7) to TG3 in the presence of IgG1 mAbs targeting epitope groups 1, 2 or 3. The mAbs were either added individually or in combination, and the degree of inhibition was calculated based on the signals obtained in the absence of competing mAb.

TG2 and TG3 have virtually identical tertiary structures comprising four domains: an N-terminal β-sandwich domain, a catalytic core domain and two C-terminal β-barrel domains.^[20]^ Most TG2-specific antibodies in CeD were shown to recognize epitopes located in the N-terminal domain or spanning the N-terminal and core domains.^[12, 21]^ To see if we could assign TG3 epitope locations, we assessed mAb reactivity to a chimeric TG3/TG2 enzyme consisting of the N-terminal domain of TG3 attached to the core and C-terminal domains of TG2. As previously observed, this protein preserves conformational epitopes and is catalytically active.^[14]^ Most of the mAbs did not recognize this chimeric protein, indicating that their epitopes include residues outside of the N-terminal domain (Figure 3C-D). However, three out of four mAbs assigned to epitope group 2 retained binding to the TG3/TG2 chimera, demonstrating that these mAbs target epitopes confined within the N-terminal domain of TG3. To understand if the epitope groups we define here reflect anti-TG3 serum reactivity in DH patients, we next assessed the ability of IgG mAbs representing epitope groups 1, 2 and 3 to block binding of serum IgA (Figure 3E). In five of seven patient sera, the epitope 3 mAb gave the highest level of inhibition, indicating that the epitope recognized by *IGHV2-5:IGKV4-1* antibodies is a major target of anti-TG3 serum IgA. Further, by combining mAbs representing all three epitope groups, we observed, on average, 40% reduction in IgA binding. Thus, antibodies targeting these epitopes constitute a substantial fraction of the total anti-TG3 reactivity in DH patients.

### TG3-specific serum IgA show biased usage of heavy and light chain V-gene segments

To further characterize the connection between serum IgA and gut plasma cells, we purified TG2- and TG3-specific IgA from serum samples of DH patients and analyzed the fractions by LC-MS/MS. Both the TG2- and TG3-specific IgA antibodies were almost exclusively of the IgA1 subclass, and both types of antibodies mostly used kappa light chains (Figure 4A-B). By assessing the distribution between individual *IGHV* and *IGKV* families, we observed biased usage of *IGHV5* and *IGKV1* segments among the TG2-specific antibodies (Figure 4C-D and S3), similar to what has previously been described for TG2-specific antibodies in CeD.^[10, 22]^ The TG3-specific antibodies, on the other hand, showed overrepresentation of *IGHV2* and *IGKV4* families. In particular, the V-gene segments *IGHV2-5* and *IGKV4-1* were more abundant among the TG3-specific antibodies compared to non-TG3-non-TG2-binding serum IgA antibodies (Figure 4E). Thus, the same V-gene segments that appeared to dominate among TG3-specific gut plasma cells were also overrepresented in TG3-specific serum IgA, indicating a close connection between systemic and mucosal anti-TG3 antibodies. Collectively, our results suggest that B cells using *IGHV2-5:IGKV4-1* play a prominent role in the anti-TG3 response in DH and that they are selectively activated to become IgA-secreting plasma cells.

**Figure 4.**
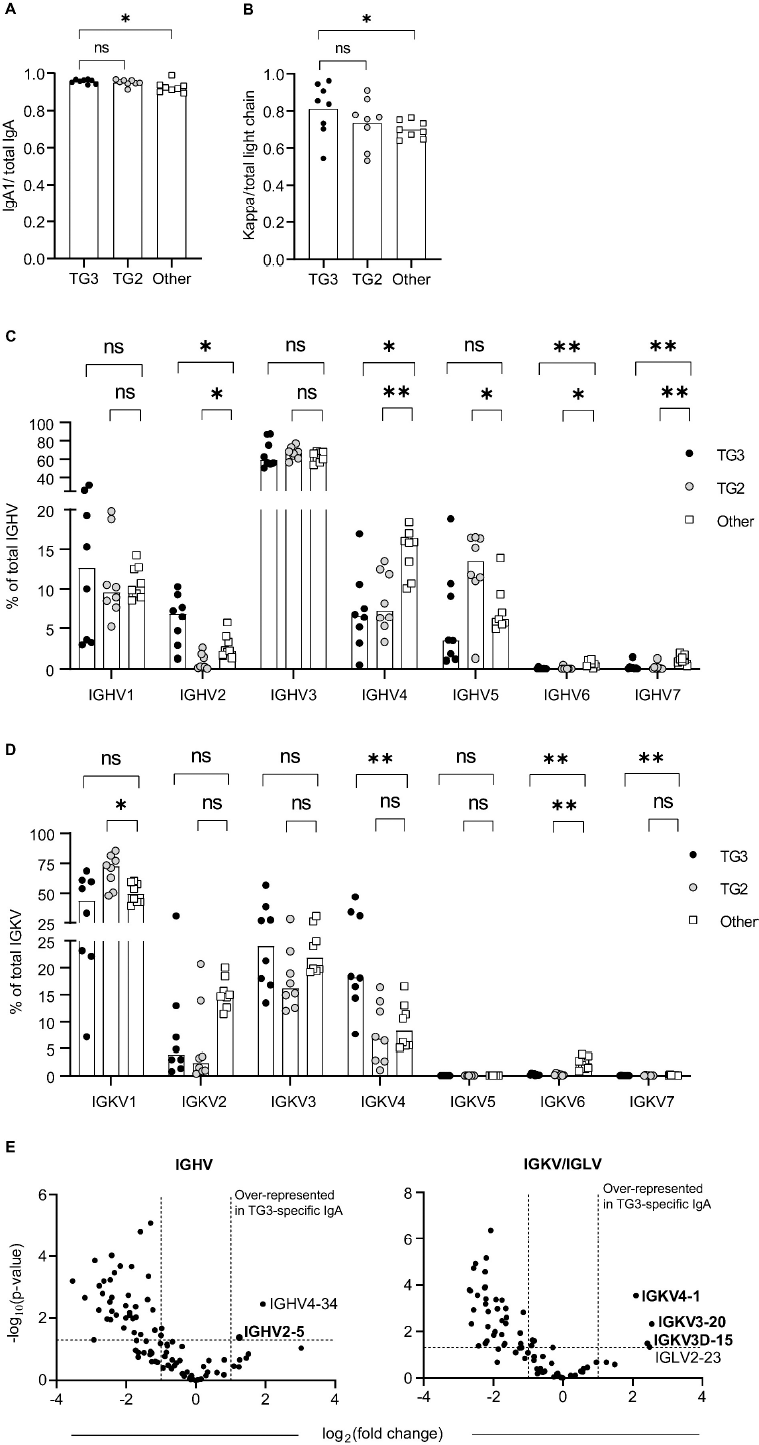
Mass spectrometry analysis of TG2- and TG3-specific serum IgA. (A-B) Relative levels of IgA1 (A) and kappa light chains (B) among TG3-specific, TG2-specific and non-TG3-non-TG2-specific (other) serum IgA antibodies isolated from sera of individual DH patients (*n*=8). Ratios were calculated from intensity-based absolute quantification (iBAQ) values obtained from MaxQuant, and difference between groups was evaluated by a paired t-test. (C-D) Distribution of the antibodies between individual *IGHV* (C) and *IGKV* (D) gene families based on iBAQ values. (E) Volcano plots showing difference in usage of individual heavy and light chain V-gene segments between TG3-specific IgA and other IgA antibodies in serum samples of DH patients. Statistical difference was evaluated by an unpaired t-test. Dashed lines indicate a p-value of 0.05 and a two-fold change in protein level based on label-free quantification (LFQ) intensity values obtained from MaxQuant. Gene segments showing overrepresentation among TG3-specific antibodies are indicated with their names. The figure shows data from four DH patients that were analyzed in one LC-MS/MS run. V-gene segments that were confirmed to be associated with TG3-specific antibodies in a second LC-MS/MS run of three DH patients are indicated in bold. **p<0.01, *p<0.05

## Discussion

The autoantibody response to TG3 is central in the pathogenesis of DH, but very little is known about its molecular details. Here, we have characterized TG3-specific autoantibodies by using antigen to isolate single plasma cells of the small intestine followed by sequencing of their rearranged Ig genes and generation of recombinant mAbs.

This study has several important take home messages: 1) The anti-transglutaminase response in DH comprises BCRs/antibodies that are specific for either TG3 or TG2 with no sign of cross-reactivity between the two. 2) All DH patients have TG2- and TG3-specific plasma cells in the gut lamina propria, with TG2-specific plasma cells usually showing higher frequency than TG3-specific plasma cells. 3) TG3-reactive BCRs/antibodies recognize conformational epitopes which can be categorized into at least three epitope groups. 4) Similar to TG2-reactive plasma cells, TG3-reactive plasma cells are newly generated CD19+ cells that have few Ig mutations compared to other gut plasma cells. 5) There is evidence for preferential usage of *IGHV2-5:IGKV4-1* by BCRs/antibodies reactive with TG3, mirroring the preferential usage of certain heavy and light chain V-gene combinations observed for BCRs/antibodies reactive with TG2.^[10, 13]^

All the DH patients studied here have prominent B-cell responses to TG2, and the fact that DH patients have T cells reactive with deamidated gluten peptides,^[4E]^ lead us to hypothesize that DH patients, as regular CeD patients, develop an immune response to gluten involving gluten-TG2 complexes. Such complexes facilitate interactions between TG2-specific B cells and gluten-specific CD4+ T cells, leading to generation of TG2-specific plasma cells as well as clonal amplification of long-lived gluten-specific T cells.^[23]^ TG3-specific antibodies are likely formed through a parallel mechanism involving gluten-specific T cells and TG3-gluten complexes formed in DH patients under conditions that are not known. In support of this mechanism, recombinant TG3 was previously observed to deamidate and, thus, form enzyme-substrate complexes with gluten peptides harboring known T-cell epitopes.^[8]^

While all DH patients had serum IgA antibodies to both TG2 and TG3, none of the tested CeD patients showed reactivity to TG3. Since DH generally develops later in life than CeD, it is likely that DH represents a late-stage complication of CeD and that the anti-TG2 response precedes the anti-TG3 response. In this regard, it would be interesting to see if detection of anti-TG3 IgA in CeD patients can predict forthcoming development of DH. In a few DH patients, there was more IgA reactivity against TG3 than TG2, but in most cases the anti-TG2 response dominated both in serum and among gut plasma cells. Collectively, our assessment of IgA reactivity in CeD and DH suggest that gluten primarily drives an anti-TG2 response and that the anti-TG3 response arises as a secondary event in some individuals, possibly through shared involvement of gluten-specific T cells in production of both types of autoantibodies.

The fact that both TG2-specific and TG3-specific plasma cells have fewer Ig mutations than other IgA gut plasma cells supports the existence of parallel mechanisms in their development. Our results give no credence to the notion that, in DH patients, TG3-reactive antibodies have developed from TG2-reactive antibodies. Conceivably, clonal amplification of gluten-specific CD4+ T cells through interactions with TG2-specific B cells is required for formation of TG3-specific plasma cells. Importantly, our characterization of TG3-specific BCRs should allow in future experiments to test whether hapten-carrier-like complexes of TG3 and gluten will allow crosstalk between TG3-specific B cells and gluten-specific T cells.

Another question that needs to be answered in future work is where pathogenic TG3 is located. Expression of TG3 has been demonstrated in hair follicles, esophagus, epidermis, brain, stomach, spleen, small intestine, testes, and skeletal muscles.^[24]^ In mice, TG3 is abundantly expressed in the colon,^[25]^ where it contributes to the stabilization of the mucus layer by crosslinking of mucin-2.^[26]^ Whether TG3 exerts a similar function in humans is not known. Notably we observed little or no expression of TG3 in human colon, but we observed expression of TG3 in the crypt compartment of the small intestine. By contrast, we observed abundant expression of TG2 expression both in the small intestine and the colon. The low intestinal expression of TG3 compared to TG2 could, at least in part, explain why the B-cell response to TG2 dominates over that to TG3 in most DH patients.

The expression pattern of TG3 in the gut is relevant for the phenotype of TG3-specific plasma cells, as plasma cells in the small intestine and colon represent clonally distinct populations of cells, with cells primed at inductive sites of the small intestine primarily producing IgA1 and cells primed at inductive sites of the colon showing skewing toward IgA2.^[27]^ Our observation that TG3-specific IgA in serum of DH patients is primarily of the IgA1 subclass indicates that the immune response leading to their formation has taken place at immune inductive sites of the upper gastrointestinal tract and not the colon. Further, the finding that the ratio between TG2- and TG3-specific gut plasma cells reflects the relative serum antibody levels suggests that the small intestine is equally relevant for the two antibody responses.

The results provided here give support to the hypothesis that formation of autoantibodies to TG2 and TG3 follow the same modus operandi. A third type of anti-transglutaminase autoantibodies targeting TG6 have been implicated in gluten-related neurological disorders.^[28]^ Future work should address if these are also generated by a similar mechanism. Collectively, our results demonstrate that detailed characterization of antigen-specific BCRs/antibodies indeed can provide useful information for deciphering the pathogenesis of autoimmune conditions.

## Materials and methods

### Patient material

Duodenal biopsies and peripheral blood samples were collected from DH and CeD patients, who were on a gluten-containing diet, as well as from control donors. The diagnosis of DH and CeD were made according to established guidelines.^[29]^ All participants gave informed consent. Biological samples from 22 DH patients were collected at Tampere University Hospital with ethical approvals provided by the Regional Ethics Committee of Tampere University Hospital, Finland. In 14 out of the 22 DH patients (ethical approval R7042), samples (serum from all subjects, gut biopsies from 4 subjects) were collected at the time of the diagnosis. From the remaining 8 patients (ethical approval R16039), serum was obtained after an oral gluten-challenge at the time of the disease relapse (median of 3.5 months) as described previously^[30]^. Out of the 22 DH patients, 68% were male, and the median age was 51 (range 23-77) years. At the time of the study, 19 out of 22 had small bowel villous atrophy, 20 out of 22 were positive for serum TG2-antibodies, and granular IgA deposits were detected in the dermis by direct immunofluorescence in 19 out of 21 DH patients with available samples. In addition, biological material was collected at Oslo University Hospital with ethical approvals from the Regional Committee for Medical Research Ethics South-East Norway. We analyzed serum from 14 CeD patients and 14 control subjects (ethical approval REK ID 6544), and we performed immunofluorescence staining of TG3 and TG2 on skin tissue from one healthy donor (ethical approval REK ID 11040) and unaffected ileum and colon tissue from two cancer patients (ethical approval REK ID 9080).

### Production of recombinant human transglutaminases

Recombinant human transglutaminases were expressed in Sf+ insect cells using baculovirus as previously described.^[31]^ Briefly, cDNA encoding human wild-type TG2 or TG3 with an N-terminal His6-tag followed by a BirA recognition sequence was cloned into the pAcAB3 vector harboring the *Eschericia coli* biotin-protein ligase gene (BirA). A chimeric TG3/TG2 protein consisting of the N-terminal domain of TG3 (aa 1-136) followed by the core and C-terminal domains of TG2 (aa 141-687) was produced in the same way. Addition of D-biotin (5 µM) to the expression cultures resulted in complete site-specific biotinylation of the expressed proteins.^[32]^ The proteins were purified from cell lysates by Ni-NTA affinity chromatography followed by buffer exchange into Tris-HCl (20 mM, pH 7.4), NaCl (300 mM), EDTA (1 mM), DTT (1 mM).

### Flow cytometry and single-cell sorting

Duodenal biopsies were processed into single-cell suspensions by treatment with collagenase followed by cryopreservation as previously described.^[13]^ In order to identify antigen-specific plasma cells, biotinylated TG2 and TG3 were attached to APC-conjugated streptavidin (Agilent) and PE-conjugated streptavidin (Thermo), respectively, using a molar antigen:streptavidin ratio of 1.5:1. Unoccupied binding sites were blocked with free D-biotin before the labeled antigens were added to the single-cell suspensions together with the following antibodies: Anti-human CD38-FITC (clone HB7, Thermo), anti-human CD19-PerCP/Cy5.5 (clone HIB19, BioLegend), anti-human IgA-APC/Vio770 (clone IS11-8E10, Miltenyi), anti-human CD3-Brilliant Violet 510 (clone OKT3, BioLegend), anti-human CD14-Brilliant Violet 510 (clone M5E2, BioLegend). Dead cells were labeled with LIVE/DEAD fixable Aqua (Thermo). Live TG3-binding IgA plasma cells (identified as large CD38^hiIgA+CD3-CD14-^ lymphocytes) were single-cell sorted into 5 µl 1% (v/v) Nonidet P40 Substitute, Tris-HCl (20 mM, pH 8.0) containing 5 U murine RNase inhibitor (NEB) using a FACSAriaIII instrument (BD). The cell lysates were flash-frozen on dry ice and stored at −80°C until further use.

### Generation of recombinant mAbs

Heavy and light chain variable regions were amplified from single-cell cDNA by a nested PCR approach as previously described.^[33]^ The PCR products were cloned into expression vectors for production of full-length human IgG1 molecules in HEK293-F cells followed by purification of the secreted mAbs on Protein G columns (GE Healthcare). Sequence analysis was done with the IMGT/V-QUEST tool.^[34]^

### ELISA

Recombinant human TG2 and TG3 proteins were coated at 3 µg/ml in Tris-buffered saline (TBS) followed by incubation with different concentrations of mAbs or serum in 0.1% (v/v) Tween-20/TBS (TBST) containing 3% (w/v) BSA. Bound IgA or IgG was detected using alkaline phosphatase-conjugated goat anti-human IgA (Sigma) or goat anti-human IgG (Southern Biotech) antibodies. To assess the dependence of antibody binding on correct TG3 conformation, the enzyme was denatured with urea (8 M) for 30 min prior to coating. Equal coating levels of native and denatured TG3 were confirmed by detection of the biotinylated proteins with alkaline phosphatase-conjugated streptavidin (Southern Biotech).

### Competition ELISA

Competition for TG3 binding was assessed using unlabeled and biotinylated mAbs. Each mAb was biotinylated by incubation with 1 mM Sulfo-NHS-LC-biotin (Thermo) in PBS followed by removal of free biotin using Zeba Spin desalting columns (Thermo). For each biotinylated mAb, a concentration falling within the linear range of the ELISA was used for incubation with coated unbiotinylated TG3 (Zedira) in the presence or absence of a 100-fold excess of unlabeled competitor mAb. Binding of the biotinylated mAb was subsequently detected using alkaline phosphatase-conjugated streptavidin (Southern Biotech). The ability of IgG1 mAbs to compete with serum IgA for TG3 binding was assessed in a similar way. Coated TG3 was incubated in the presence or absence of 25 µg/ml of each mAb before addition of purified serum IgA (see below) to reach a concentration falling in the linear range of the ELISA. Bound IgA was subsequently detected as described above.

### Adsorption of TG2- and TG3-binding serum antibodies

To test for serum antibody cross-reactivity, serum samples were incubated with 20 µl M-280 Streptavidin Dynabeads (Thermo) that had been pre-coated with 2 µg biotinylated TG2 or TG3 in TBS. The tubes were placed in a magnetic separator, and IgA reactivity to TG2 and TG3 in the supernatant was assessed by ELISA as described above.

### Purification of serum antibodies and mass spectrometric analysis

TG2- and TG3-specific IgA was purified from ∼1 ml of DH sera and analyzed by liquid chromatography tandem mass spectrometry (LC-MS/MS) essentially as previously described.^[22]^ Briefly, total IgA purified on Peptide M-agarose (Invivogen) was incubated with M-280 Streptavidin Dynabeads coated with 10 µg biotinylated TG2 or TG3. The non-binding fraction was collected by magnetic separation, and the beads were washed extensively before TG2- and TG3-specific antibodies were eluted with glycine-HCl (0.1 M, pH 2.5). The eluted antibodies were immediately neutralized with Tris-HCl (1 M, pH 9.0) and denatured by addition of urea (8 M) in Tris-HCl (100 mM, pH 8.0), before they were reduced with DTT and alkylated with iodoacetamide. The samples were then diluted with Tris-HCl (100 mM, pH 8.0) and digested with 0.5 µg sequencing-grade trypsin (Promega). The generated peptides were desalted and analyzed in duplicates by LC-MS/MS using a timsTOF fleX mass spectrometer equipped with a nanoElute LC system (Bruker). The data were searched against a database containing the amino acid sequences of all human heavy and light chain V-gene segments obtained from the International ImMunoGeneTicS Information System (https://www.imgt.org/) using MaxQuant software, version 2.0.3.0.^[35]^

### Microscopic analysis

Snap-frozen tissue biopsies of histologically normal human skin, ileum and colon was cut at 4 μm and sections were adhered to SuperFrost slides by thaw-mounting and air-dried. Sections were stained with rabbit-anti-human TG2 (custom made polyclonal antibody from Pacific Immunology, CA) or with rabbit-anti-human TG3 (PA596450, Thermo Fischer Scientific) followed by detection with Alexa 488 donkey anti-rabbit (Molecular Probes) antibody. Slides were counterstained with 4’,6-diamidino-2-phenylindole (DAPI) and mounted with ProLong Diamond Antifade Mountant (ThermoFisher). Images were acquired using a 20X objective lens Zeiss LSM 880 Confocal Laser Scanning Microscope.

### Statistical analysis

GraphPad Prism version 9.3.1 was used to perform statistical analysis. Centers are given as means, and difference between groups was evaluated by paired or unpaired two-tailed t-tests unless otherwise indicated in the figure legends. p values <0.05 were considered significant.

## Acknowledgements

We thank Bjørg Simonsen and Marie K. Johannesen for excellent technical assistance. Anna Alakoski, Noora Nilsson, and Eriika Mansikka at the Celiac Disease Research Center, Tampere University and Department of Dermatology, Tampere University Hospital, are thanked for partaking in patient examinations. Flow cytometry and cell sorting experiments were performed at the Flow Cytometry Core Facility, Oslo University Hospital. We are also thankful to Maria Stensland and Sachin Singh at the Proteomics Core Facility for conducting mass spectrometry analyses. The work was supported by grants from Stiftelsen KG Jebsen (project SKGJ-MED-017), the University of Oslo World-leading research program on human immunology (WL-IMMUNOLOGY), the South-Eastern Norway Regional Health Authority (project 2020027), the Academy of Finland (projects 347471 and 347473), the Sigrid Juselius Foundation (project 8069) and the Competitive State Research Financing of the Expert Responsibility area of Tampere University Hospital (9AB068 and 9AC088).

## Author Contributions

Conceptualization: Saykat Das, Rasmus Iversen, Ludvig M. Sollid; Funding Acquisition: Ludvig M. Sollid, Katri Lindfors, Teea Salmi; Patient examination and sample collection: Esko Kemppainen, Kaisa Hervonen, Katri Lindfors, Teea Salmi, Knut E.A. Lundin, Jørgen Jahnsen; Investigation: Saykat Das, Jorunn Stamnaes, Rasmus Iversen; Naveen Parmar; Frode L. Jahnsen Supervision: Rasmus Iversen, Ludvig M. Sollid; Visualization: Saykat Das, Rasmus Iversen, Ludvig M. Sollid; Writing – Original draft: Saykat Das, Rasmus Iversen, Ludvig M. Sollid. Writing – Review and Editing: All authors.

---

**Figure S1.**
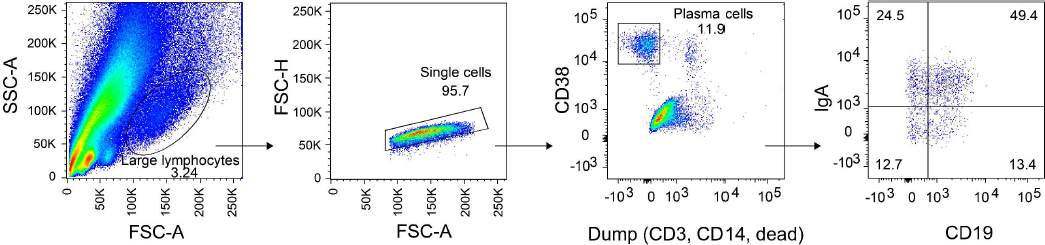
Gating strategy. Representative flow cytometry plots showing identification of IgA plasma cells in duodenal biopsy single-cell suspensions of DH patients.

**Figure S2.**
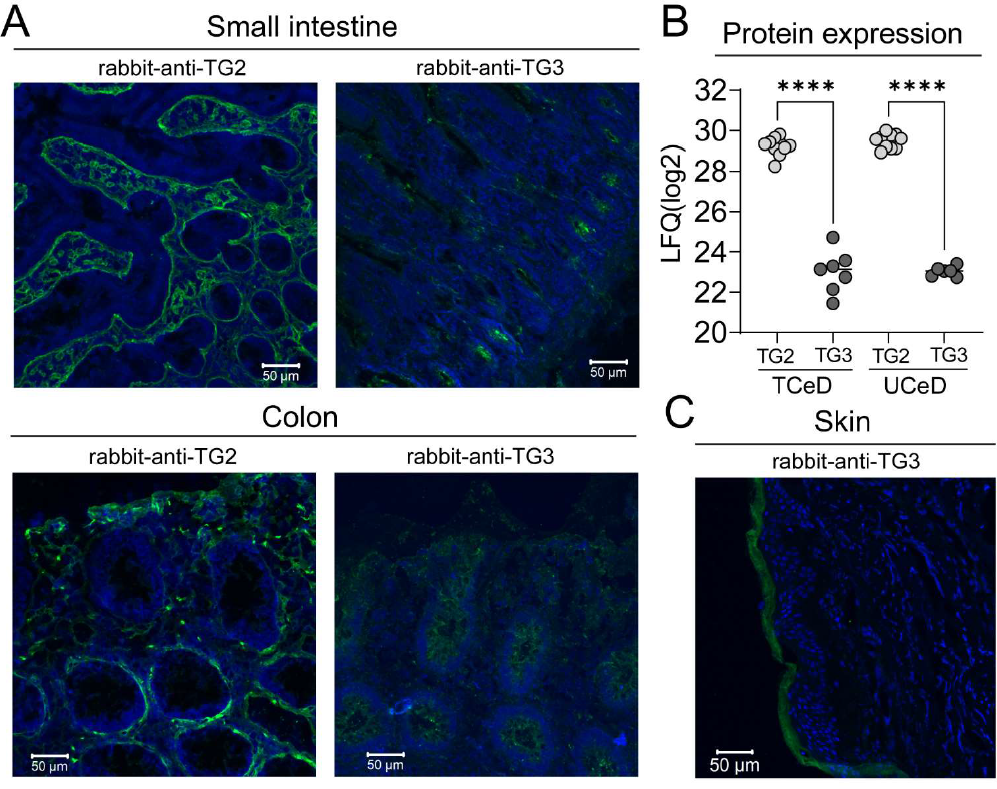
TG2 and TG3 protein expression. (A) Staining pattern for TG2 (polyclonal rabbit anti-human TG2 antibody) and TG3 (polyclonal rabbit anti-human TG3 antibody) in unfixed frozen sections from human small intestine and colon. Nuclei were stained with 4′,6-diamidino-2-phenylindole (DAPI). Scale bars; 50µm. (B) Comparison of TG2 and TG3 protein expression in human small intestine. The plot shows expression values from a previously published dataset of LC-MS/MS based proteome analysis of FFPE biopsy tissue sections from treated (TCeD) and untreated (UCeD) patients.^[19]^ Proteins were quantified by label-free quantification (LFQ). Each circle represents expression values from one patient biopsy block. Expression was compared using one-way ANOVA with Tukey’s adjustment for multiple testing (****p<0.0001). (C) Immunofluorescence staining of human skin using polyclonal rabbit anti-human TG3 antibody (green). Positive TG3 staining is observed in stratum corneum of epidermis.

**Figure S3.**
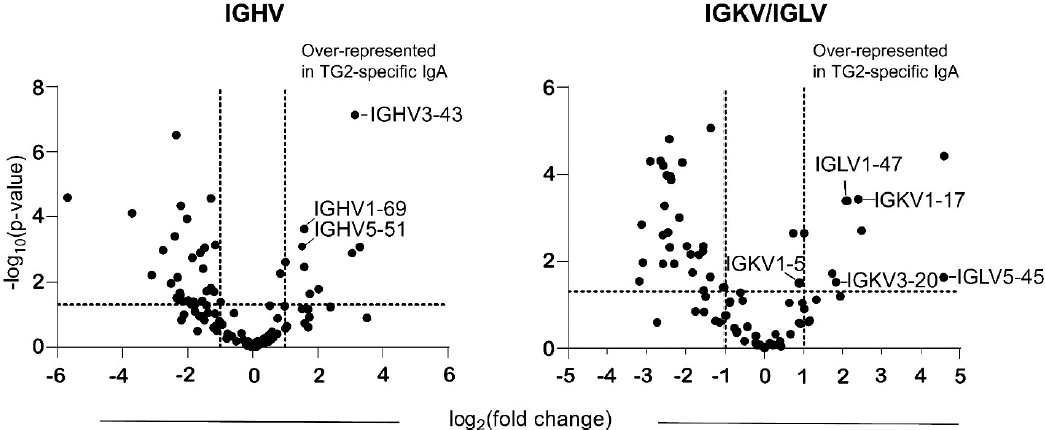
V-gene usage by TG2-specific serum IgA in DH patients. Volcano plots showing difference in *IGHV* and *IGKV/IGLV* usage between TG2-specific IgA and other IgA antibodies isolated from serum samples of DH patients (*n*=4). Statistical difference was evaluated by an unpaired t-test. Dashed lines indicate a p*-*value of 0.05 and a two-fold change in protein level based on label-free quantification (LFQ) intensity values obtained from MaxQuant. V-gene segments previously observed to be overrepresented among TG2-specific antibodies in CeD are indicated with their names.

**Table S1.**
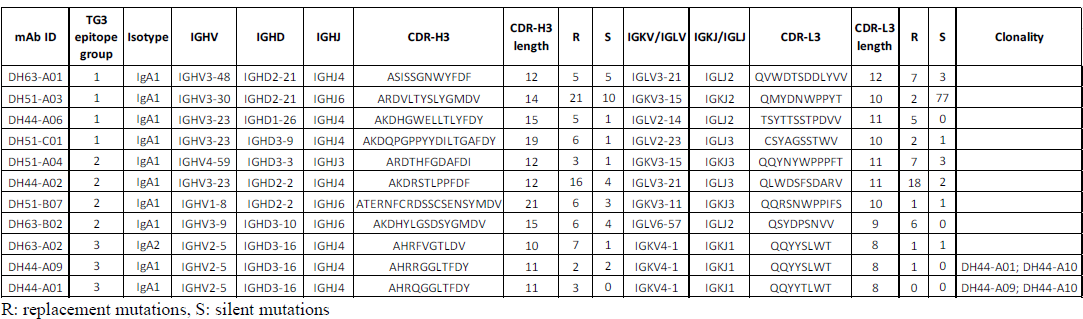
Sequences and sequence-properties of TG3-specific mAbs generated from gut plasma cells of DH patients

